# Chemo-mechanical Coupling in the Transport Cycle of a Type II ABC Transporter

**DOI:** 10.1101/471920

**Authors:** Koichi Tamura, Hiroshi Sugimoto, Yoshitsugu Shiro, Yuji Sugita

## Abstract

AT P -binding cassette (ABC) transporters are integral membrane proteins that translocate a wide range of substrates across biological membranes, harnessing free energy from the binding and hydrolysis of ATP. To understand the mechanism of the inward- to outward-facing transition that could be achieved by tight regulation of ATPase activity through extensive conformational changes of the protein, we applied template-based iterative all-atom molecular dynamics (MD) simulation to the heme ABC transporter BhuUV-T. The simulations, together with biased MDs, predict two new conformations of the protein, namely, occluded (Occ) and outward-facing (OF) conformations. The comparison between the inward-facing crystal structure and the predicted two structures shows atomic details of the gating motions at the transmembrane helices and dimerization of the nucleotide-binding domains (NBDs). The MD simulations further reveal a novel role of the ABC signature motifs (LSGG[Q/E]) at the NBDs in decelerating ATPase activity in the Occ form through sporadic flipping of the side chains of the LSGG[Q/E] catalytic serine residues. The orientational changes are coupled to loose NBD dimerization in the Occ state, whereas they are blocked in the OF form where the NBDs are tightly dimerized. The chemo-mechanical coupling mechanism may apply to other types of ABC transporters having the conserved LSGG[Q/E] signature motifs.

## Introduction

Chemo-mechanical couplings where large conformational changes of a protein are coupled to chemical events (i.e., binding and hydrolysis of nucleotide triphosphates) are often found in biomolecular motors,^1–3^ signaling proteins,^3^ chaperones,^4^ and clock proteins.^5^ The existence of the coupling implies that appropriate positioning^6^ of the catalytically competent residues is realized only in a catalytic-dwell state of a protein and is disrupted in other conformational states of the cycle to prevent futile usage of ATP. For example, the formation of the catalytic-dwell state of the myosin motor is coupled to a global conformational change that brings the catalytic loop (switch II) closer to the ATP-binding P-loop.^7,8^ Elucidation of chemo-mechanical coupling mechanism is central to understanding biomolecular machines including ATP-binding cassette (ABC) transporter.

ABC transporters are integral membrane proteins that are ubiquitous in all life forms and mediate the translocation of diverse substrates across the lipid bilayer, harnessing the free energy gained by the binding and hydrolysis of ATP.^9–11^ They are involved in a vast variety of biological processes and their malfunction due to mutation often leads to serious medical conditions such as cystic fibrosis^12^ and multidrug resistance.^13^ Much attention has been paid to their transport mechanisms with a view to developing effective therapies.

ABC transporters have a common architecture: two transmembrane domains (TMDs) that form a substrate binding pocket, and two nucleotide-binding domains (NBDs) that bind and hydrolyze ATP (Figure 1a). Some ABC transporters have covalently linked TMD and NBD, while in others the domains are separate. The TMDs and the NBDs are physically connected by coupling helices (CHs) which transmit the conformational change at the NBDs to the TMDs. Each NBD has a well-conserved^10,11^ phosphate-binding motif (P-loop)^14^ and an ABC signature motif (LSGG[Q/E]).^15^ It is well established that the binding of ATP induces dimerization of the NBDs in a head-to-tail fashion in which the P-loop of one monomer sandwiches ATP with the signature motif of the opposite monomer (Figure 1b).^16,17^ The dimerization realizes the formation of competent catalytic sites where the side chains of opposing monomers are oriented toward the substrate so that the transition state(s) of the hydrolysis reaction is stabilized.

**Figure 1.**
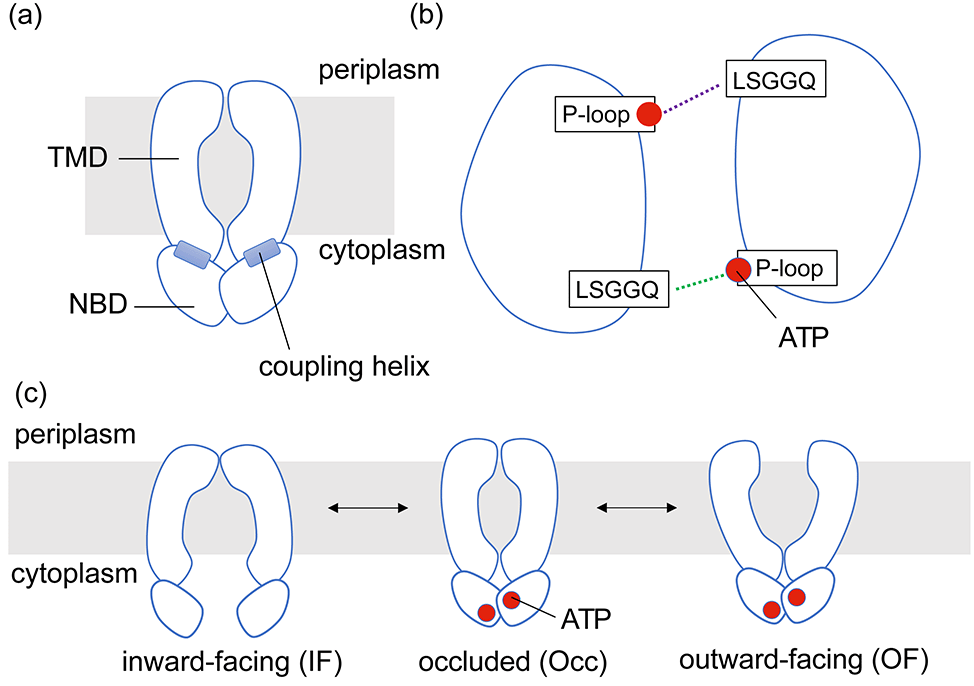
Architecture and alternating access mechanism of ABC transporter. (a) A shared architecture of ABC transporter. Transmembrane domains (TMDs) are embedded in the biological membrane, whereas nucleotide-binding domains (NBDs) are exposed to the cytoplasm. They are physically connected by the coupling helices. (b) Dimerization interface of NBDs as seen from the periplasmic side. They dimerize upon binding of ATP to the P-loop in a head-to-tail fashion, where the P-loop of one monomer binds to the LSGG[E/Q] motif of the opposing monomer. (c) An alternating access mechanism of ABC transporter. The conformational transition from the inward-facing (IF) to the outward-facing (OF) form involves the binding of ATP to the NBDs and the formation of the occluded (Occ) state.

The translocation of substrates by ABC transporters is thought to occur by an alternating access mechanism (Figure 1c).^10,11^ In this mechanism, the structure of the ABC transporter alternates between inward-facing (IF) and outward-facing (OF) forms. The formation of an occluded (Occ) intermediate during the conformational transition between IF and OF makes the substrate binding pocket inaccessible to bulk solvent and prevents reverse transport. This is a key feature of transporters, and contrasts with channel proteins.

ABC transporters are classified into exporters and importers according to the substrate transport direction.^9^ Importers, which are found only in prokaryotes, are further classified into type I, type II and energy coupling factor (ECF) transporters.^10,11^ Each class of importer has a different substrate transport strategy and a specific structural fold. Type I and II importers are associated with a periplasmic binding protein (PBP) that binds substrate in the periplasm and delivers it to the TMDs of the ABC transporter.^10,11^ Type I importers transport relatively small compounds such as sugars, while type II importers are specific for the transport of trace elements.^10,11^ The ECF transporters share a unique substrate acquisition mechanism and are devoted to the transport of micronutrients.^18^

Atomic structures of full ABC transporters largely come from experimental studies using X-ray crystallography^10,11^ and/or cryo-electron microscopy.^19^ These techniques, however, often yield the protein structure of only one of the possible conformational states and relatively unstable intermediates along the functional cycle are difficult to elucidate, which is a major obstacle to the understanding of the coupling mechanism. In this respect, atomistic molecular dynamics (MD) simulations are a promising approach to reveal experimentally inaccessible conformational states in transporters.^20–30^ Homology (comparative) modeling techniques^31,32^ are also widely applied to predict elusive conformational states in a protein.^33,34^ Since these techniques do not take into account thermal motions of proteins, the conformational stability of putative structures needs to be carefully examined by MD simulation.

In this study, we employ template-based iterative atomistic MD simulation to predict hitherto unknown conformations of heme importer of *B. cenocepacia* (BhuUV-T) ^35^ and provide insights into the mechanism of the IF-to-OF transition occurring after substrate translocation. BhuUV-T belongs to the type II ABC importer family and transports heme (Fe-porphyrin complex) across the inner membrane of bacteria. Heme importers in pathogenic bacteria play a major role in the acquisition of iron, which is utilized during infection and propagation. BhuUV-T is a dimer of a dimer (BhuU (TMD) + BhuV (NBD)), together with a PBP (BhuT) (Figure 2a and S1). The coupling helices (CHs), which physically link the NBDs and the TMDs, are located between TM6 and 7 (Figure 2a and S1). BhuUV-T has been crystalized in an apo IF conformation with and without BhuT (throughout this manuscript, we use the term “apo” for the structure without bound nucleotides and substrates) and their structures have been characterized crystallographically.^35^ The apo IF conformation with bound BhuT (Figure 2a) may represent a post-translocation state, in which a heme has already been released towards the cytoplasm. The transport cycle is postulated to proceed via the binding of ATP at the NBDs (BhuVs) to eventually form the OF state. The formation of the OF state would be accompanied by full dimerization of the NBDs (BhuVs) and the dissociation of BhuT. As mentioned, the cycle for transporters likely involves the transient formation of an Occ state after the translocation of substrate. Formation of the Occ state and closure of the cytoplasmic gate are thought to result from the binding of ATP and dimerization of the NBDs (BhuVs) (Figure 1c). The question which then arises is: how is the hydrolysis of ATP prevented in the Occ state, such that futile cycling back to the IF state is avoided, before formation of the OF state? To solve this problem, we analyzed MD trajectories of BhuUV-T starting from predicted Occ and OF conformations and arrived at a novel chemo-mechanical coupling mechanism that regulates ATPase activity by way of reorientation of the serine residues of the LSGG[Q/E] motifs in type II ABC transporters. The proposed mechanism likely extends to other types of ABC transporters, as dimerization of NBDs is a common feature.^10,11^

**Figure 2.**
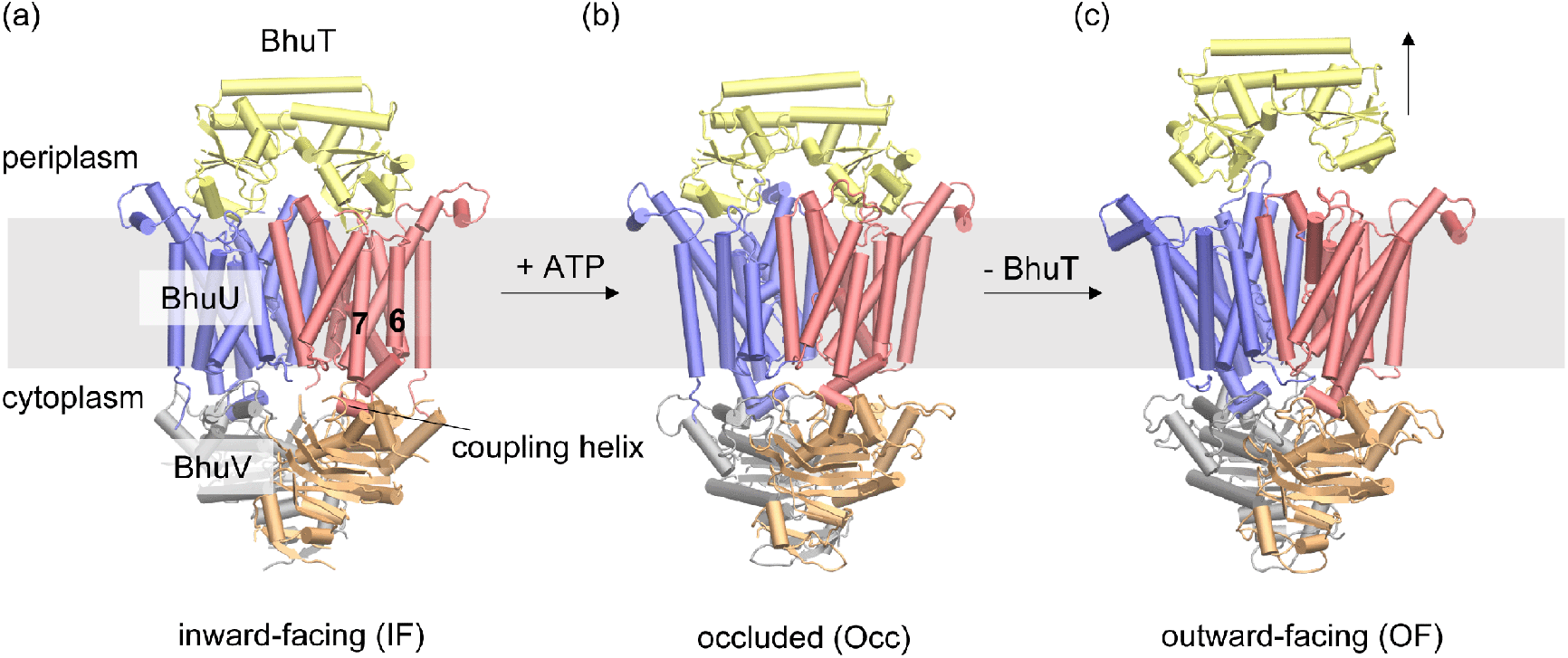
Structures of heme importer BhuUV-T. (a) Crystal structure of BhuUV-T complex (PDB 5B58) in the inward-facing conformation. Transmembrane domains (TMDs, dimer of BhuU) are shown in blue or red. Nucleotide-binding domains (NBDs, dimer of BhuV) are shown in gray or orange. Periplasmic binding domain (PBP, BhuT) is shown in yellow. (b,c) Alternate conformations of BhuUV-T predicted in this study. (b) The occluded and (c) the outward-facing conformations.

## Computational Methods

### Overall strategy

In this study, we focus on the IF-to-OF transition of heme importer BhuUV-T (Figure 2). Accordingly, our simulation systems do not contain the heme substrate. The unknown OF structure of the heme importer (Figure 2c) was modeled by template-based iterative MD simulation. The technique consists of iterative refinements of rough model by all atom MD simulations which consider solvent and membrane environment. The initial rough model was generated with homology modeling^31,32^ using the structure of the vitamin B12 transport protein BtuCD in the ATP analog-bound form as template (PDB 4R9U).^36^ Since 2002 several crystal structures of type II ABC importers have been elucidated, but the BtuCD structure is the only OF structure with bound nucleotides (Table S1). The engineered disulfide bond introduced in the BtuCD structure to suppress basal ATPase activity was not included in the homology model. The relatively low sequence identity (~34%) between BhuUV and BtuCD poses a significant challenge for homology modeling of thermally stable structures in the membrane environment. We therefore extensively refined the predicted structure by a series of MD simulations. Modeling procedures are briefly summarized in the following section, and details are described in SI (Text S1). The structural stabilities of the predicted structures were examined by relatively long (~1 μs) MD simulations without any restraints on protein atoms.

We first considered the nucleotide-bound Occ crystal structure of BtuCD-F (PDB 4FI3)^37^ as a template of the Occ form, however, the lower resolution (3.47 Å) compared to that of the OF form (2.79 Å) could have introduced further uncertainties into the modeling process. Instead, the Occ form (Figure 2b) was modeled by targeted MD (tMD) simulation^38^ using part of the modeled OF structure as target (see below). The effect of the binding of ATP to the NBDs was then investigated by comparing the predicted structures with the crystal structure of the apo IF form (Figure 2). Representative MD simulations are summarized in Table S2.

### Iterative refinements of the OF form

We employed MODELLER version 9.16 for building the homology models.^31,32^ Sequence alignments were generated by the align2d() function in MODELLER, using the template 3D structure to put gaps in the alignment. For example, the function avoids placing gaps within the secondary structure. Altogether, we generated four structural models (MODEL1–4). The first OF model with bound ATP, which was based on an alignment involving several manual adjustments (MODEL1), turned out to be unstable after a short MD simulation of ~20 ns: we observed spontaneous closure of the periplasmic gate and distortion of the helices at Arg121 to Gly134 of both NBD monomers (Figure S2a and S2b). The instability, especially in the helical region in the NBDs, seemed to originate from the rather low sequence identity between BhuUV and the vitamin B12 transporter BtuCD used as template (Figure S2c). On the other hand, there is a crystal structure of a heme importer of *Y. pestis* (HmuUV) in the apo OF form (PDB 4G1U)^39^ having a higher sequence identity (~40%) which can generate better models. Note that the *apo* HmuUV structure has a significantly different structural arrangement compared to that of the *nucleotide-bound* BtuCD and therefore cannot be used as a template for the modeling of the *nucleotide-bound* OF form of the *B. cenocepacia* heme importer BhuUV. As an alternative, we generated a reference homology model (REF1) by using the apo HmuUV structure as template and replaced the unstable regions in MODEL1 with the corresponding ones of the REF1 structure to make a chimera. Although the helical regions in the new model (MODEL2) turned out to be stable, some other parts were still unstable: the periplasmic gate again spontaneously closed and there was disruption of the dimer interface (data not shown). We speculated that the structural instability may have arisen from the low sequence identity of TM6 connected to the CHs (Figure S1) enlarging the dimer interface by way of the structural coupling. Then, we built the next model (MODEL3), which involved replacing TM6 in MODEL2 with that in the REF1 structure. Although MODEL3 was stable during the first ~600 ns of the MD simulation, the periplasmic gate gradually closed after this and eventually closed almost completely (Figure S3a). We also observed fraying in the helical region at Ala51 to Ala70 in one of the BhuU monomers (Figure S3b). The gradual closure of the periplasmic gate involved TM5a of one monomer leaning towards the other (Figure S3a). Then, the final model, MODEL4, was generated by replacing the fraying region and the leaning helix of MODEL3 with the corresponding parts of the opposing monomer. This model turned out to be stable during 1.5 μs MD simulation (MD-2ATP-OF) and, importantly, the periplasmic gate stayed open (Figure S4a and S4b). Ca root mean squared deviation (RMSD) between the template (PDB 4R9U) and the average structure of the predicted OF form calculated from the last 500-ns part of the MD trajectory was 2.2 Å (Figure S4c). More details of the modeling are found in SI.

### Modeling of the Occ intermediate

Targeted MD (tMD) simulation^38^ starting from the equilibrated ATP-bound IF structure was employed to model an Occ conformation of BhuUV-T with bound nucleotides. tMD has been widely used to elucidate the conformational transition pathways of proteins including a type II ABC transporter.^40^ In tMD, a biasing force is applied to pull the protein towards a predefined target structure. The application of the external force enables one to induce conformational changes in proteins very efficiently. A caveat is that this method is known to be subject to “large-scales-first” bias and the pathway may deviate from the actual physical one.^41,42^ We thus disregarded the pathway itself and checked only for stability in the final model. Note that if one chooses the entire MODEL4 OF structure as a target of tMD, the simulation would end up with generating the target structure itself. Alternatively, if one selects a part of the MODEL4 structure as a target, one would obtain protein with the selected part having conformation of the MODEL4 and the remaining part left almost unaffected. In our case, the NBDs and the residues of the cytoplasmic gate of the equilibrated ATP-bound IF structure were pulled so that the domains and the gate closed as in the MODEL4 OF structure (see SI for details). We then performed an equilibrium MD simulation without the bias potential on the protein atoms to check the structural stability of the predicted Occ form. However, this first model was unstable: there was immediate opening of the cytoplasmic gate and widening of the distances between coupling helices in a short MD simulation (data not shown). Then, a second model was generated by also pulling TM6 and 7 of both monomers towards the target (MODEL4). An equilibrium MD simulation (1.5 μs) confirmed the structural stability of the predicted Occ form (MD-2ATP-Occ, Figure S5a). Ca-RMSD between the averaged structure along the last 500-ns part of the MD trajectory of the Occ form and the BtuCD-F Occ crystallographic one was 2.5 Å (Figure S5b).

### MD Simulation details

All MD simulations were performed with the development version of GENESIS.^43,44^ The force field parameter set for water molecules was TIP3P,^45,46^ and those for protein and lipids were CHARMM36.^47,48^ We used revised parameters for ATP^49^ and magnesium ions.^50^ The protein was embedded in a 1-palmitoyl-2-oleoyl-sn-glycero-3-phosphoethanolamine (POPE) bilayer. Typical simulation system size was 134 × 134 × 197 Å^3^ (~360,000 atoms) for the BhuUV-T system and 134 × 134 × 154 Å^3^ (~280,000 atoms) for the BhuUV system. The equation of motion was integrated with the velocity Verlet algorithm with a time step of 2.5 fs. Bonds including a hydrogen atom were constrained by the SHAKE/RATTLE^51^ (non-water molecules) and the SETTLE^52^ (water molecules) algorithms. Temperature and pressure were regulated by the stochastic velocity rescaling thermostat^53^ and the barostat,^54^ respectively. The thermostat and the semi-isotropically coupled barostat were applied every 10 steps. Long range electrostatic interactions were calculated with the particle mesh Ewald method^55^ and updated every 2 steps. Short range Lennard-Jones interactions were cutoff at 12 Å with a force switching function beginning at 10 Å.^56^ Details are described in SI. MD trajectories related with the manuscript are available at this link https://osf.io/hfb2c/.

## Results

### Characteristics of the Predicted Occ and OF Structures

Crystal structures of heme importer BhuUV and vitamin B12 transporter BtuCD have identified two gates for heme transport, i.e., the cytoplasmic gate and the periplasmic gate. The cytoplasmic gate, especially the so-called cytoplasmic gate II, is formed by Asn108 and Leu110 (numbering of BhuUV) of opposing monomers which are well conserved among type II ABC transporters (Figure 3a).^37^ The periplasmic gate of BhuUV(-T) is formed by Asp200 and Arg204 of opposite subunits which are conserved among heme importers (Figure 4, left).^35^ In the IF state of the BhuUV-T crystal structures, the cytoplasmic gate II is widely opened, while the periplasmic gate is closed through a salt bridge between Asp200 and Arg204 (Figure 3a and Figure 4, left).^35^ Eventually, the PBP BhuT stably associates to the periplasmic surface of TMD of BhuUV through the electrostatic interaction of Glu94 or Glu231 (BhuT) and Arg84 (BhuU) (Figure S6).^35^ In the MD simulations of the IF and Occ states, however, those salt-bridges were disrupted, and instead salt-bridge pairs between Glu94 or Glu231 and Arg81 (Bhu) were formed (Figure S5c and S6).

**Figure 3.**
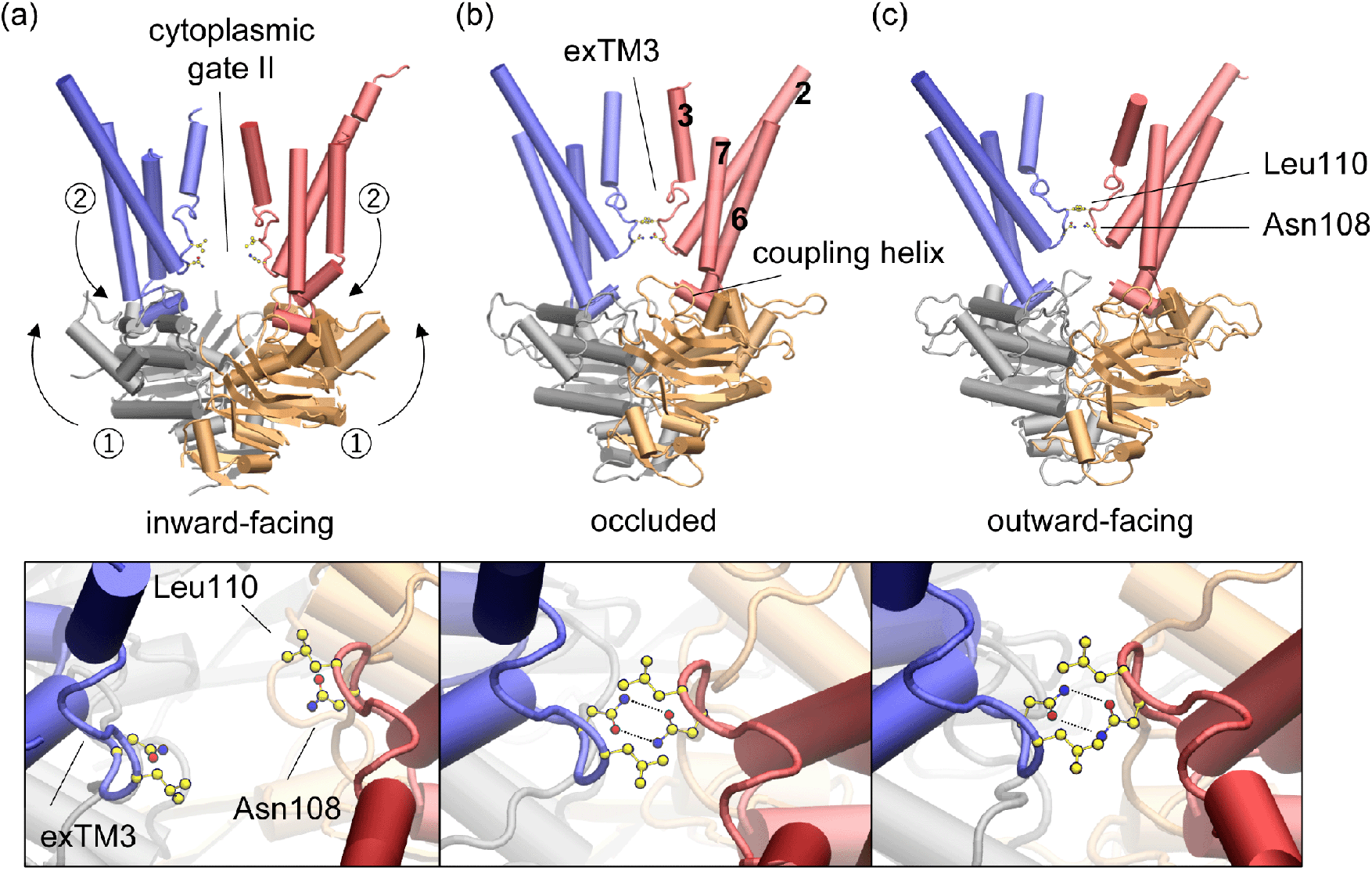
Closure of the cytoplasmic gate II coupled to NBD dimerization. The cytoplasmic gate is open in the inward-facing (IF) conformation (a), while it is closed in the occluded (Occ) (b) and the outward-facing (OF) (c) conformations. The upper panels show the transporter from the membrane space and the lower ones from the periplasmic space. Only TM2–3 and 6–7 are shown for the TMDs. Carbon, nitrogen, and oxygen atoms are in yellow, blue, and red, respectively.

**Figure 4.**
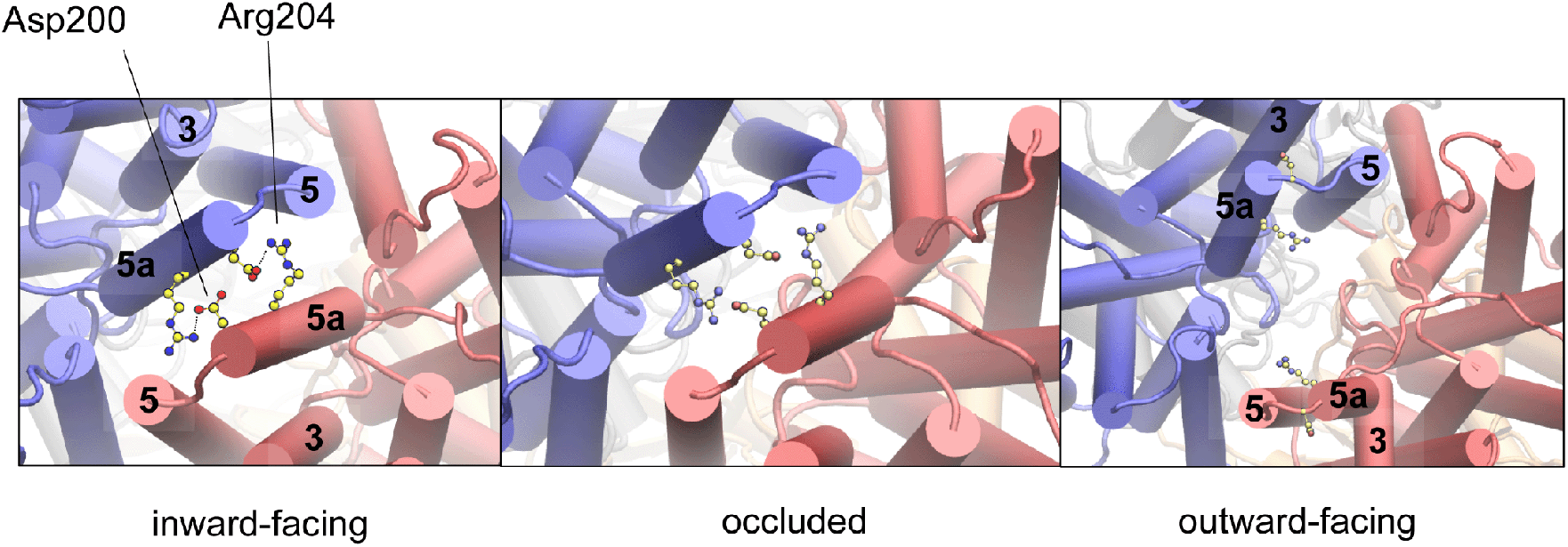
Opening of the periplasmic gate upon the binding of ATP. The panels show the transporter from the periplasm space. In the inward-facing (IF) state (left, PDB 5B58), the periplasmic gate is sealed by TM5 and 5a of opposing monomers. The inter-domain salt bridges between Asp200 and Arg204 stabilize the dimer interface. The situation is not changed in the predicted occluded (Occ) state (middle), while in the predicted outward-facing (OF) state (right), the gate is wide open due to the large tilt of the TM helices.

In our predicted Occ structures of BhuUV-T, both gates are closed and BhuT still binds to TMD (Figure 3b and 4, middle). In addition, the central cavity, although large enough to accommodate the substrate heme, is filled with water molecules (Figure S7a). The predicted Occ structural characteristics fit well with the crystal structure of the Occ state of the BtuCD-F complex with bound ATP analogs (PDB 4FI3), in which a large central cavity for substrate is also observed (Figure S7b).^37^ On the other hand, in the predicted OF structure of BhuUV, the periplasimic gate is opened, while the cytoplasmic gate II is closed (Figure 3c and 4, right). Details of the conformational change from IF to the unique Occ conformation found in our simulation are described in the following section.

Differences between the predicted OF and Occ forms of BhuUV(-T) and the crystal structures of BtuCD (PDB 4R9U)^36^ and BtuCD-F (PDB 4FI3)^37^ are best characterized by the distance between the coupling helices (CHs). The distance in both predicted structures is shorter (~2–4 Å) than that in the corresponding crystal structures (Figure 5). A shorter distance causes a greater tilt of TM6 and 7 of opposing monomers (Figure S8), and the strong tilt of the TM helices, especially in the OF form, stabilizes an open periplasmic gate. However, in the MD simulation of the ATP-bound OF BtuCD with the engineered disulfide bond (BtuCD-2ATP-OF) there is spontaneous closure of the periplasmic gate within a short period (Figure S9). Spontaneous closure was often observed in the simulation of the apo OF BtuCD (PDB 1L7V) whose periplasmic gate is structurally similar to that of the ATP-bound form (PDB 4R9U).^20,57,58^ In contrast, the simulation of the predicted OF form of BhuUV finds a state in which the periplasmic gate is stably opened due to a greater tilt of the TM helices in the membrane environment (Figure S4b). Therefore, the modeled structures of BhuUV-T are likely more relevant to physiological conditions, while the BtuCD(-F) crystal structures seem trapped in slightly unnatural states possibly due to the introduction of the artificial disulfide bond and the mutations. Indeed, the mutated BtuCD proteins did not exhibit ATP hydrolysis activity.^37^

**Figure 5.**
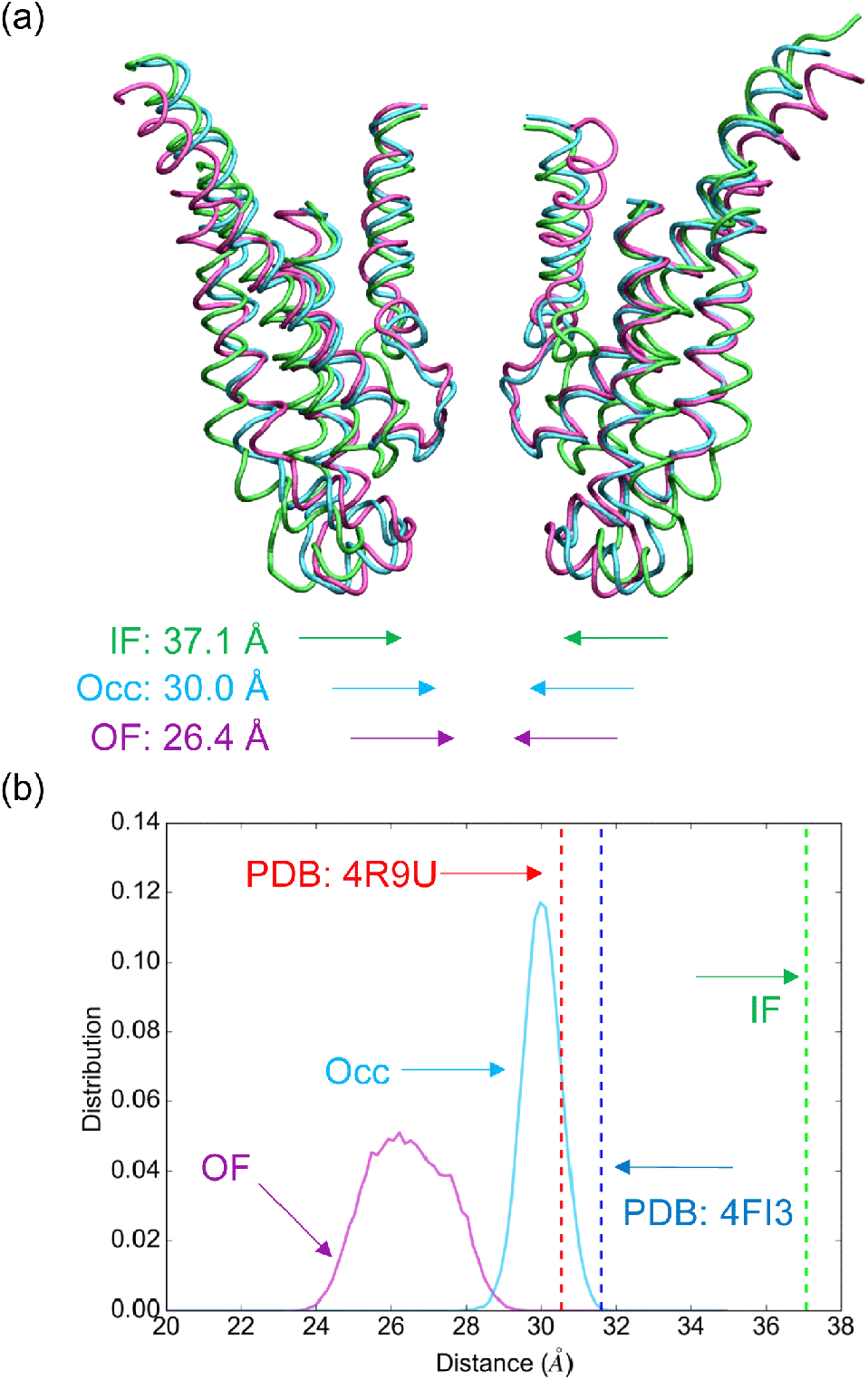
Comparison of the distance between the coupling helices of type II ABC importers in different conformational states. (a) The predicted OF (magenta) and Occ (sky blue) average structures along the last 500-ns equilibrium MD simulations are superimposed on the IF crystal structure (green). Only TM2–3 and TM6–7 of the TMDs are shown. Distance between the coupling helices (CHs) for each structure is denoted at the bottom. (b) Distribution of distance between the CHs in the last 500-ns equilibrium MD simulation for each conformation.

### NBD Dimerization Induces Closure of Cytoplasmic Gate and Opening of Periplasmic gate

The opened cytoplasmic gate II in the IF structure is due directly to separation of the extended stretch (exTM3) of opposing monomers (Figure 3a). In the initial step from the IF state to the Occ state, the binding of ATP promotes dimerization of the NBDs (Figure 3a, ①), and eventually forces the coupling helices (CHs) and the TM6/7 bundles to lean towards the central cavity (Figure 3a, ②). These conformational changes pushes TM2, and forces its C-terminal part to tilt towards the central cavity, eventually closing the cytoplasmic gate II bordered by the exTM3s (Figure 3b). Tight closure of the gate was maintained throughout the MD simulations of the Occ state (Figure S5d) and of the OF state (Figure S4d).

The disposition of the periplasmic gate of BhuUV(-T) in the IF crystal structure persists in the predicted Occ state despite large conformational changes in the NBDs and several TM helices (Figure 4, middle, and Figure S5e). On the other hand, in the OF state, the periplasmic gate is opened due to the tilt of TM5 and 5a together with other helices (Figure 4, right, and Figure S4b). The large tilt of the helices on the periplasmic side of the TMD is coupled to the dissociation of BhuT from TMD BhuU, as discussed later. Gate opening and closing is highly correlated with dimerization of the NBD. We measured the distance between two CHs from each NBD monomer for the IF, Occ and OF structures. As shown in Figure 5, the average distance is 37.1, 30.0 and 26.4 Å, in the MD simulation of the IF, Occ and OF states, respectively. The dimerized forms of Occ and OF states are somewhat different. The inter-NBD distance of the Occ form gradually increases and deviated from that of the OF form (Figure S10). We defined the NBD structure of Occ and OF as “partial” and “full” dimerization forms, respectively.

### Stoichiometry of ATP in the IF-to-OF transition of BhuUV-T

The role of bound ATP in closing the cytoplasmic gate II was examined in a series of MD simulations. In the predicted Occ and OF forms of BhuUV(-T), two ATPs are tightly bound to the active sites and contribute to the tight closure of cytoplasmic gate II (Figure 3). This is similar to what is observed in the corresponding crystal structures of BtuCD(-F).^36,37^ Consistently, when we added two ATPs to the NBDs of the apo IF BhuUV-T complex, the CHs, which directly contact the NBDs, tended move closer (Figure 6a). The importance of ATPs is further highlighted by the following observation: when we removed two ATPs from the predicted Occ state, there was spontaneous widening of the CH distance (Figure 6a). Importantly, the narrowing and widening of the CH distance clearly correlates with the tendency to formation and destabilization of the cytoplasmic gate II, respectively (Figure 6b), demonstrating mechanical coupling between the NBD and the TMD. Destabilization of the dimer interface upon removal of ATPs is also seen in the BtuCD-F system.^59^

**Figure 6.**
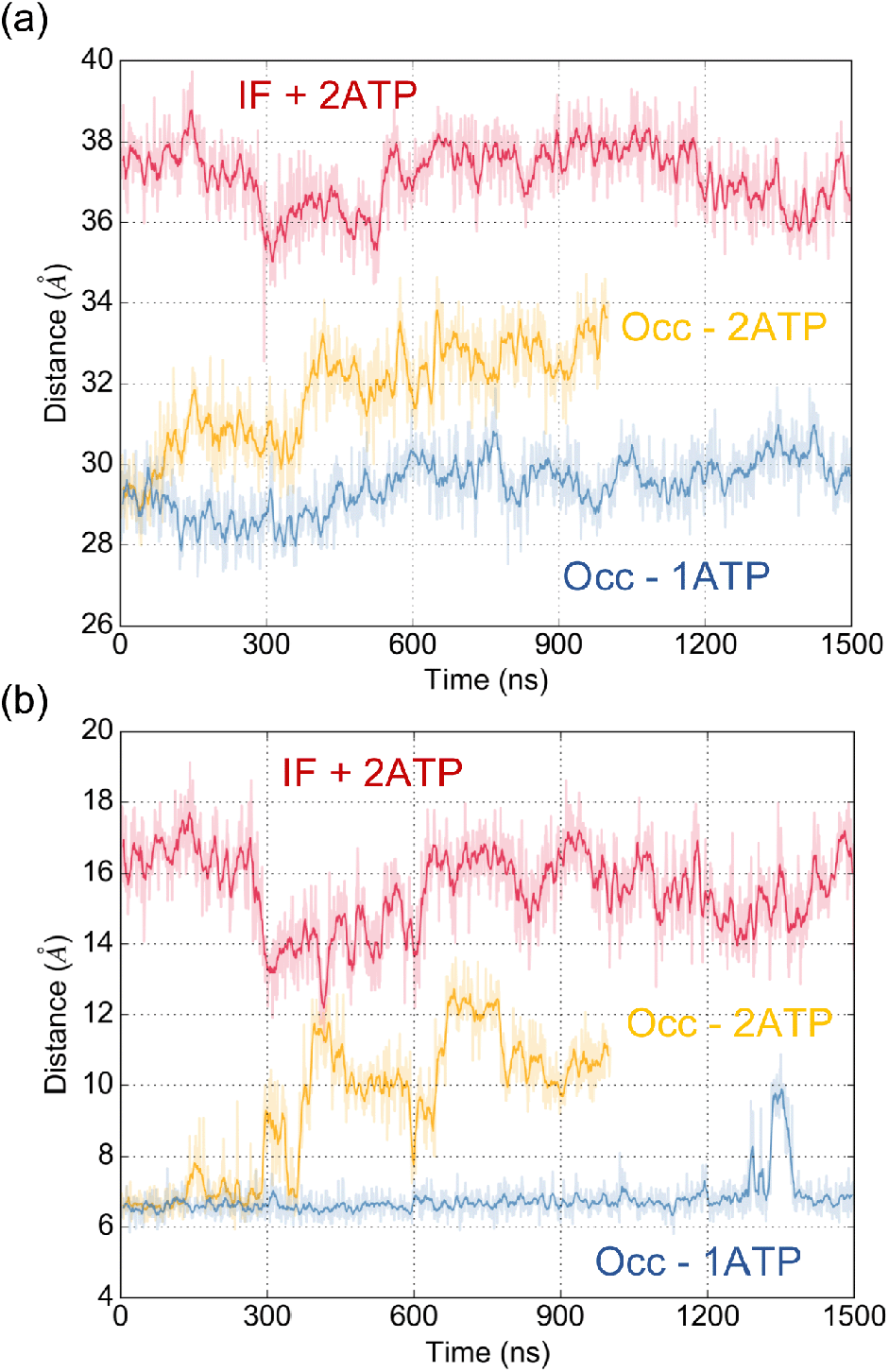
Structural changes in the equilibrium MD simulations. (a,b) Temporal changes of distance between (a) the coupling helices and (b) Leu110 of opposing monomers during the MD-IF-2ATP (IF + 2ATP), MD-apo-Occ (Occ – 2ATP), and MD-Occ-1ATP (Occ – 1 ATP) trajectories.

Is *one* ATP enough to stably close cytoplasmic gate II? To answer this, we removed one ATP from an arbitrarily chosen active site of the predicted Occ form. Although the result was not so as clear as in the simulations above, the distance between the CHs consistently widened (Figure 6a). The widening transiently opens the cytoplasmic gate (Figure 6b). These results suggest that one ATP may be insufficient to stably close the cytoplasmic gate, and a longer simulation may clarify this.

### Dissociation of the PBP from TMD Facilitates NBD Dimerization

In the MD simulation of the predicted Occ form with bound ATP, BhuT was stably bound to the TMDs, as shown by the distance between conserved^35^ salt-bridge pairs between the TMDs and BhuT (Figure S5c). The observation seems inconsistent with our previous pull-down assays performed on BhuUV-T, which show that the binding of ATP, not its hydrolysis, induces dissociation of BhuT.^35^ Therefore, the Occ form appears to be in a BhuT-dissociation dwell state with a longer dwelling time than the simulation time scale (1.5 μs).

A comparison between Occ and OF structures showed that the CHs and the helix bundles (TM6/7) in the OF form lean further than those in the Occ form, resulting in the opening the periplasmic gate (Figure 4 and 5). However, these structural changes could not be allowed in the BhuUV-T Occ form, probably due to the capping by the bound BhuT. Therefore, it was indicated that the dissociation of BhuT (PBP) from TMD of BhuUV and the structural change on the periplasmic side are coupled with each other, which is followed by “full” dimerization of the NBDs (Figure 2c). In other words, PBP (BhuT) dissociation facilitates NBD (BhuV) dimerization.

### Orientation of the Catalytic Serine Residues

During MD simulation of the Occ state, we witnessed orientational changes of the catalytically important serine residues (Ser147)^60^ in the LSGG[Q/E] motifs in the NBDs (Figure 7a, upper panel). Namely, the side chain of Ser147 in either monomer occasionally flipped between the g-phosphate of ATP and the glutamate residue of the LSGG[Q/E] motif (Figures 7a, 7c and S11a). The change in orientation from the potentially catalytic to the non-catalytic forms clearly correlates with a subtle increase in the distance between the P-loop and the LSGG[Q/E] motif (Figure 7a) which sandwich ATP (Figure 7c), due to the “partial” dimerization.

**Figure 7.**
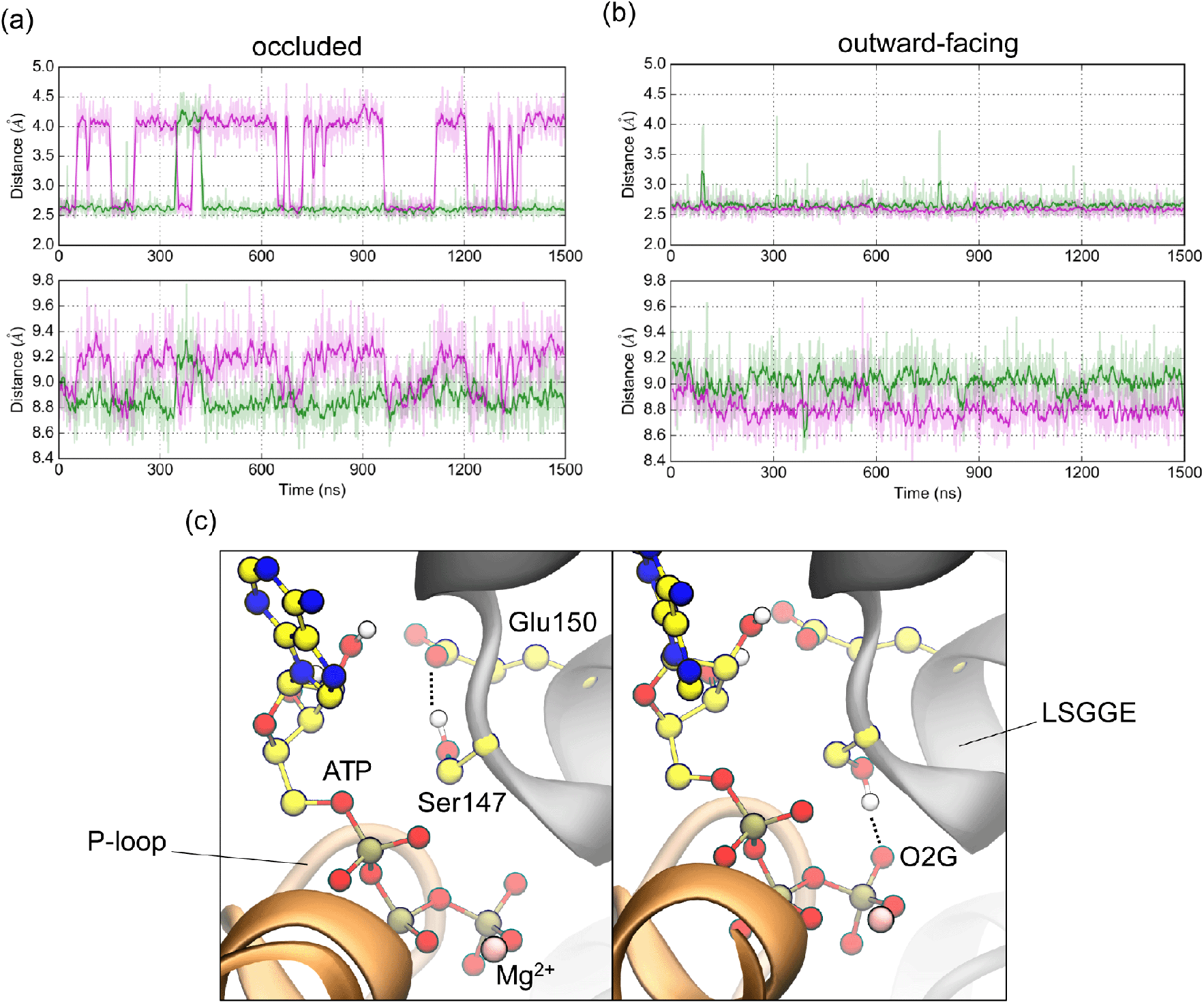
Alternate configurations of Ser147. (a,b) The upper panel shows the distance between OG atom of Ser147 and O2G atom of ATP and the lower one shows the distance between P-loop and LSGGE for the occluded state (a) and the outward-facing states (b). Here, the two nucleotide-binding sites (NBSs) are designated as NBS1 and NBS2, and the results for NBS1 and NBS2 are indicated by magenta and green lines, respectively. (c) Two possible orientations adopted by the side chain of Ser147 observed in the simulation of the occluded state.

As shown in Figure 7a, an interesting finding in the Occ state is that the side chain swiveling occurred more frequently in one of the nucleotide-binding sites than the other. Here, we designate the former as NBS1 and the latter as NBS2, respectively. The difference in side-chain mobility of Ser147 originates from the asymmetrical dimerization in the Occ state. The distance between the P-loop and the LSGG[Q/E] motif in NBS1 is larger than that in NBS2, resulting in more frequent dissociation of the side chain of the serine residue (Figure 7a, lower panel).

In contrast, in the OF state, Ser147 of both monomers is always perfectly oriented toward the γ-phosphate of ATP (Figure 7b, upper panel) presumably due to the stable “full” dimerization of the NBDs (Figure 7b, lower panel and Figure S10). The “full” dimerization (Figure S10) realizes shorter distances and holds the serine residues in a catalytic orientation (Figure 7b). Ser147 appears key in modulating the catalytic power of the protein in the functional cycle, as discussed below.

## Discussion and Conclusions

In this study, we predicted two unknown conformations of a type II heme ABC importer BhuUV-T in order to elucidate the mechanism by which the futile usage of ATP in the Occ state can be avoided. The results lead us to propose a chemo-mechanical coupling mechanism where the Ser residues (Ser147) of the ABC signature LSGG[Q/E] motifs play a fundamental role in modulating catalytic activity of the transporter (Figure 8). In this mechanism, the progress of conformational changes of the protein determines ATPase activity, in the following manner:

1. In the IF state, the protein is in an ATP-binding dwell and the catalytic sites are “empty” (without bound ATP), as represented by the crystal structure (PDB 5B58).
2. The binding of ATP to NBD (BhuV) of the IF form triggers a conformational change to the Occ form. The Occ form is apparently in a BhuT-dissociation dwell, because the bound BhuT hinders further tilting of TM helices and narrowing of the inter-CHs space. The structural constraint holds the NBDs somewhat apart due to structural coupling between the NBDs and the CHs, stabilizing “partial” dimerization. As a result, the catalytic sites are in a “loose” state, in which the Ser residues (Ser147) of LSGG[Q/E] motifs frequently flip away from the g-phosphate of ATP, and thereby catalytic activity is partially lost. The hydrolysis reaction is decelerated.
3. In going from the Occ to OF states, dimerization of the NBDs gradually proceeds, and with the dissociation of BhuT full tilting of the TM helices and complete dimerization of the NBDs are achieved. These conformational changes gradually constraint the side chains of the two Ser147s so that their orientations become increasingly catalytic and full competency and maximal ATPase activity are realized upon final dimer engagement of the NBDs. These catalytic sites are now in a “tight” state (Figure 8).

**Figure 8.**
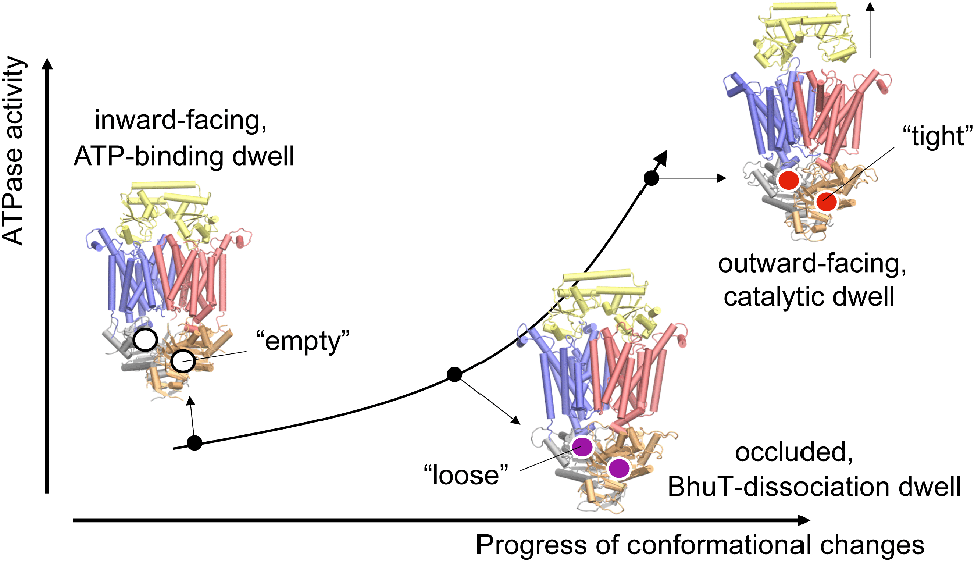
Proposed chemo-mechanical coupling model. The horizontal axis describes progress of conformational changes (e.g. distance between coupling helices), whereas the vertical axis indicates the ATPase activity of the protein. In this model, the progress of conformational changes is coupled to the ATPase activity of the protein (the curved arrow) so that as the conformation transition proceeds from the inward-facing (IF) state to the outward-facing (OF) one, the ATPase activity gradually increases.

Oldham and Chen^61^ have recently pointed out the role of the serine residues in coupling conformational change of a type I maltose ABC transporter to its catalytic activity, and that its role resembles that of the “arginine finger” of the biomolecular motor F1-ATPase, which controls the catalytic activity of the protein.^62^ However, to our knowledge, the mechanism, by which the wasteful hydrolysis of ATP in the Occ state is avoided, has not yet been addressed for any ABC transporters. Here, by combined use of structural modeling and MD simulation, the mechanism of chemo-mechanical coupling in the type II ABC transporter can be explained in atomistic detail.

With respect to the mechanism of the heme transport by BhuUV-T, our mechanism proposed for BhuUV-T is apparently similar to that for the molybdate importer MolBC, but somewhat different from those for BtuCD-F and HmuUV.^11,37,63–66^ For example, the closure of the cytoplasmic gate II in the proposed IF-to-OF transition pathway of BhuUV-T is consistent with an electron paramagnetic resonance (EPR) spectroscopy study on MolBC.^63^ Moreover, key features in the crystal structure of MolBC are consistent with those of BhuUV. Both structures have been determined in the IF form without bound ATP.^35,64^ On the other hand, the existing crystal structures of BtuCD and HmuUV without bound ATP were solved in the OF form with a closed cytoplasmic gate I, which is bordered by the N-terminal part of TM5 of opposing monomers (Table S1).

The transport cycle of a vitamin B12 transporter BtuCD-F of type II ABC importer family involves a different state that is not considered in this study.^11,37^ Namely, the ATP-binding dwell of BtuCD-F is an asymmetric apo Occ state (PDB 2QI9),^65^ unlike to BhuUV-T and MolBC (Figure S12a). Notably, the cytoplasmic side of the asymmetric apo Occ crystal structure is closed differently from BhuUV-T and MolBC. The gate of BtuCD-F in the apo Occ state is at cytoplasmic gate I. Furthermore, the transport cycle of BtuCD-F does not involve a state like the PBP-dissociation dwell (Figure S12b, middle), even though a crystal structure that could represent the state has been found for BtuCD-F (PDB 4FI3, Figure S12a). Alternatively, this latter BtuCD-F structure has been considered to be a vitamin B12-loaded Occ state despite the absence of bound substrate in the crystal structure.^11,37^ These studies imply some mechanistic divergence among the type II ABC importer family, a view supported by a recent biochemical study on HmuUV.^66^

It seems that the binding of ATP to the NBDs of the asymmetric Occ form of BtuCD-F inevitably forces dimerization of the NBDs, resulting in opening of the cytoplasmic gate I and closure of cytoplasmic gate II, as observed in the crystal structure (PDB 4FI3). Therefore, the ATP-bound form of the Occ state with the cytoplasmic gate II closed may appear between the asymmetric occluded and the OF forms, as proposed in the transport mechanism of BhuUV-T (Figure 8 and S12b). Concern over a futile hydrolysis reaction in the ATP-bound Occ form can be partially allayed by our chemo-mechanical coupling mechanism.

The dynamic nature of ABC transporters renders experimental elucidation of chemo-mechanical coupling mechanisms a challenging task. For instance, ATPase activity of BhuUV-T is measured as the time course of the concentration of released inorganic phosphate ions upon the addition of ATP.^35^ The observed kinetic parameters represent multiple dynamic processes including the IF-to-OF conformational transition and the dissociation of BhuT, complicating interpretation. The complexity behind such a single experimental parameter highlights the usefulness of our coupled computational structural modeling and MD simulation approach in deciphering chemo-mechanical coupling, as it allows for direct characterization of motions of a protein and its catalytic sites along the functional cycle. The prediction from simulation and our proposed mechanism can now be tested experimentally.

In summary, template-based iterative MD simulation and biased MD predict two structurally unknown conformations of type II heme ABC importer BhuUV-T, and provide insights into the chemo-mechanical coupling mechanism. Both structures exhibit high stability in a membrane environment and show how binding of ATP is coupled to conformational changes of the protein. The dimerization of the NBDs in the predicted Occ form is incomplete due to structural constraints imposed by bound BhuT. As a consequence, the side chains of the serine residues of the conserved LSGG[Q/E] motifs flip between catalytic and non-catalytic orientations, resulting in impaired catalytic activity. Dissociation of BhuT in the OF form enable tight dimerization of the NBDs and proper serine orientation, activating full catalytic power of the protein. These results led us to a definitive answer to the question on the futile usage of ATP in the Occ state. The proposed chemo-mechanical coupling mechanism may provide basis for the understanding of transport mechanism of many ABC transporters having different structural folds.

## Acknowledgments

K.T. thanks J. Jung, C. Kobayashi, Y. Matsunaga, and M. Kamiya for sharing their expertise on running simulations on the K computer. This work was supported by the RIKEN pioneering project “Dynamic Structural Biology” (to Y.Sugita), MEXT Grant-in-Aid for Scientific Research on Innovative Areas Grant Number 26119006 (to Y.Sugita), JP26220807 (to Y.Shiro), JP15H01655, JP17H05896 (to H.Sugimoto), and RIKEN Special Postdoctoral Researcher Program (to K.T.). This research used computational resources of the K computer provided by the RIKEN Center for Computational Science through the HPCI System Research project (to K.T., Project ID: hp170027, hp180009, to Y.Sugita, ra000009). Molecular figures were prepared with VMD^67^ and PyMOL.^68^

